## Supplementary Materials for "N-WASP-dependent branched actin polymerization attenuates B-cell receptor signaling by increasing the molecular density of receptor clusters"

**Supplementary materials for this manuscript include the following:**

7 figures

4 videos

### Figure 1-S1

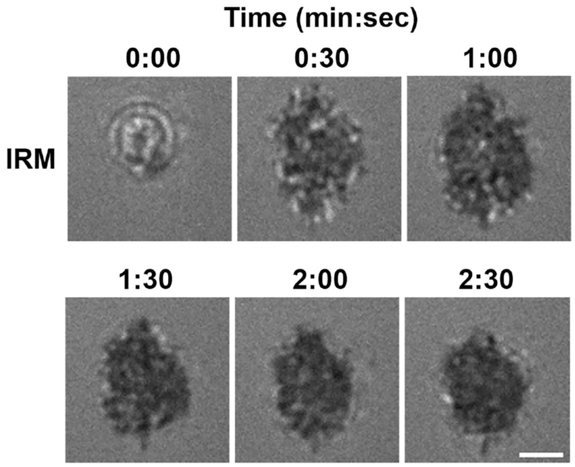


**Figure 1-figure supplement 1. B-cells spread and contract on Fab'-coated-planar lipid bilayers.** Splenic B-cells were pre-warmed to 37°C and incubated with planar lipid bilayers coated with monobiotinylated Fab' fragment of goat anti-mouse IgG+M (Fab'-PLB) and imaged live at 37°C by interference reflection microscopy (IRM). Shown are individual frames from a time-lapse IRM image of one B-cell. The plasma membrane area contacting with Fab'-PLB (B-cell contact zone) visualized by IRM increased between 0-1 min after landing, indicating spreading, and decreased after 1 min 30 sec, indicating contraction. Scale bar, 2 μm.

### Figure 1-S2

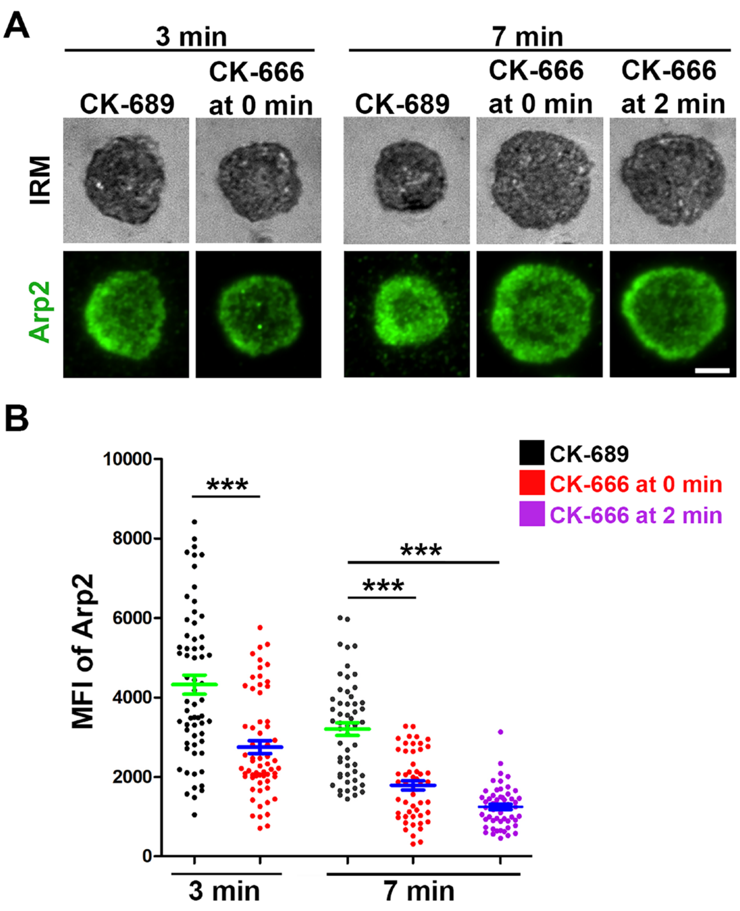


**Figure 1-figure supplement 2. CK-666 significantly decreases Arp2/3 recruitment to the B-cell contact zone.** WT splenic B-cells were treated with CK-689 or CK-666 (50 μM) before (0 min) and after maximal spreading (2 min) during incubation with Fab'-PLB at 37°C. Cells were fixed at 3 and 7 min, permeabilized, stained for Arp2, and imaged using IRM and TIRF. Shown are representative images (**A**) and the MFI of Arp2 in the contact zone at 3 min and 7 min compared between B-cell treated with CK-689 (black dots), CK-666 from 0 min (red dots), and CK-666 from 2 min (purple dots) (**B**). Data points represent individual cells from 3 independent experiments with ~20 cells per condition per experiment. Scale bar, 2 μm. *** $p$<0.001, by non-parametric student's $t$-test.

**Figure 1-S3**

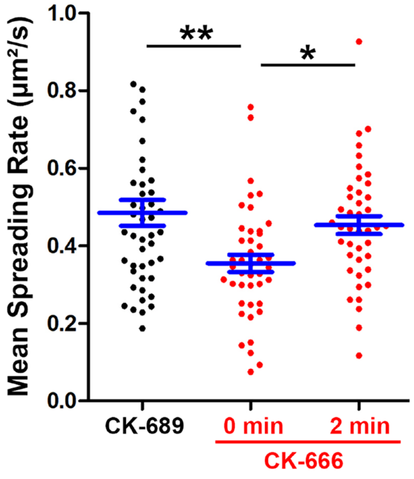


**Figure 1-figure supplement 3. CK-666 treatment before but not after maximal B-cell spreading decreased the spreading kinetics.** WT splenic B-cells were treated with CK-689 or CK-666 (50 µM) before (0 min) and after maximal spreading (2 min) during incubation with Fab'-PLB and imaged live at 37°C by IRM. The area occupied by the B-cell contact zone was measured using IRM images and custom codes made in MATLAB. The mean spreading rate of each cell during its early spreading phase was quantified using the contact area versus the time curve of that cell by linear regression. The averaged spreading rates (±SEM) were generated from 3 independent experiments with ~15 cells per condition per experiment. * *p*>0.05, ** *p*<0.01, by non-parametric student's *t*-test.

### Figure 1-S4

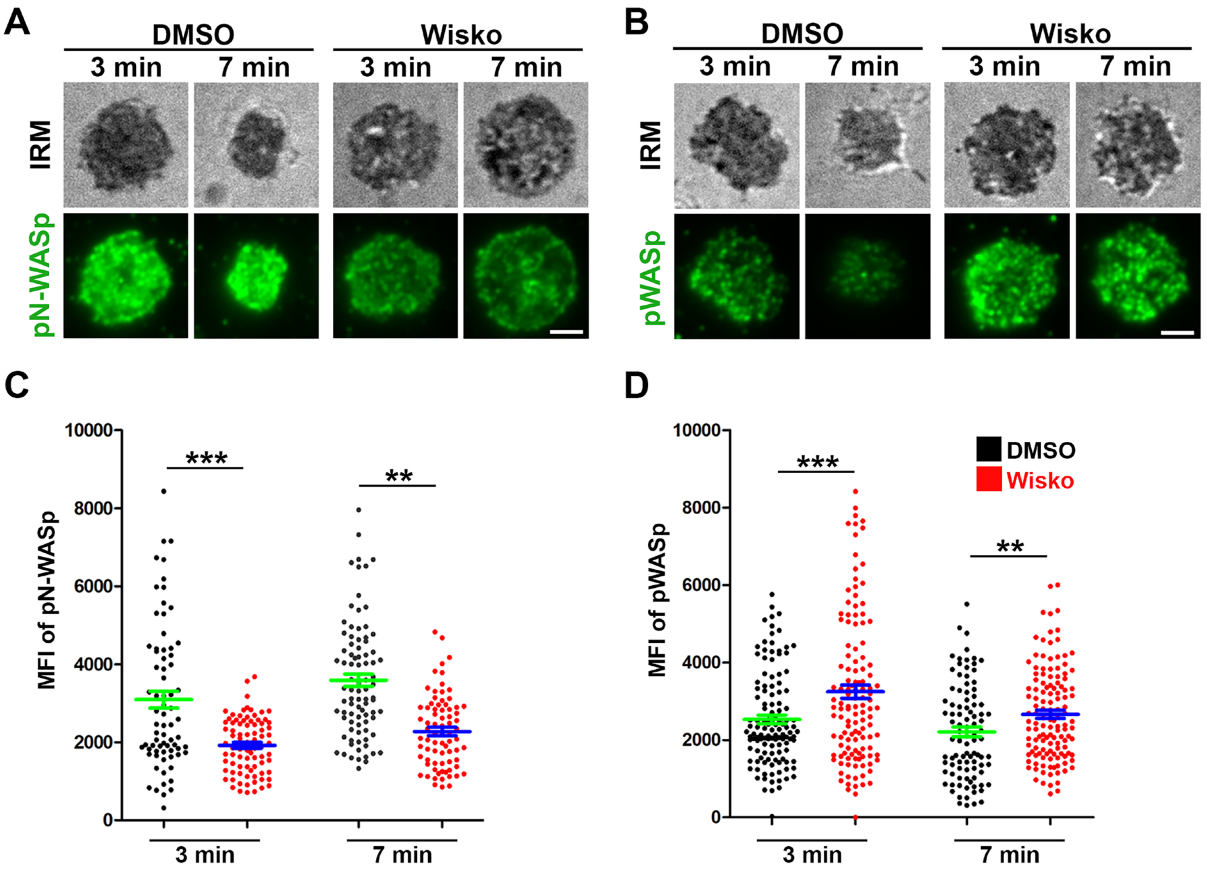


**Figure 1-figure supplement 4. Wiskostatin treatment inhibits N-WASP activation while enhancing WASP activation in B-cells.** WT splenic B-cells were pre-treated with Wiskostatin (Wisko, 10 μM) or DMSO (control) for 10 min at 37°C before and during incubation with Fab'-PLB. Cells were fixed at 3 and 7 min, permeabilized, stained for phosphorylated N-WASP (pN-WASP) or WASP (pWASP), and imaged using IRM and TIRF. Shown are representative IRM and TIRF images of pN-WASP (**A**) and pWASP (**B**) at the B-cell contact zone and the MFI (±SEM) of pN-WASP (**C**) and pWASP (**D**) in the B-cell contact zone, comparing between DMSO and Wisko-treated B-cells. Data points represent individual cells from 3 independent experiments with ~25 cells per condition per experiment. Scale bar, 2 μm. ** $p$<0.01, *** $p$<0.001, by non-parametric student's $t$-test.

### Figure 3-S1

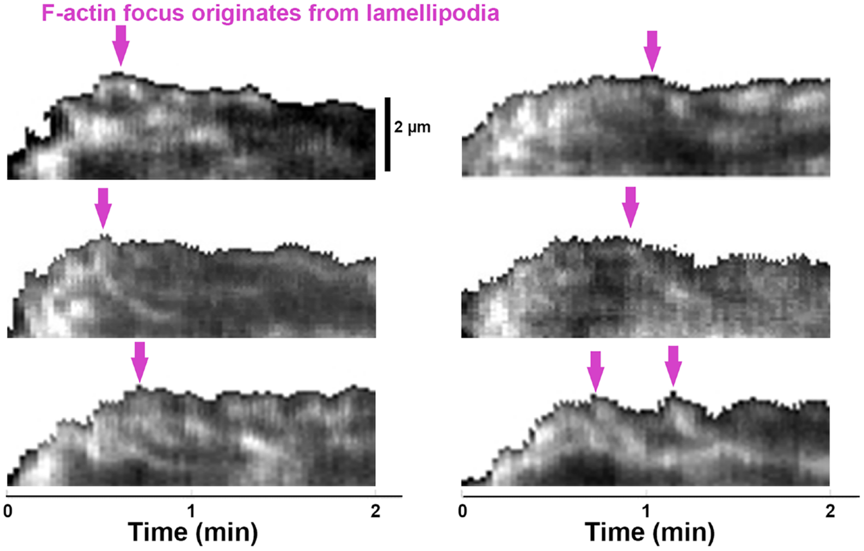


**Figure 3-figure supplement 1. Emerging of Inner F-actin foci from lamellipodia.** Splenic B-cells from LifeAct-GFP transgenic mice were treated with DMSO, imaged live using TIRF and IRM during incubation with Fab'-PLB at 37°C, and analyzed using kymographs that were randomly generated from each cell. Shown are six examples of the kymographs used for analysis. Arrows indicate the emergence of inner F-actin foci near the lamellipodia. Lamellipodia-derived inner F-actin foci were identified by their LifeAct-GFP FI ≥2 fold of their nearby region, inside location in the contact zone, migrating away the lamellipodial F-actin, and trackable for ≥8 sec.

**Figure 6-S1**

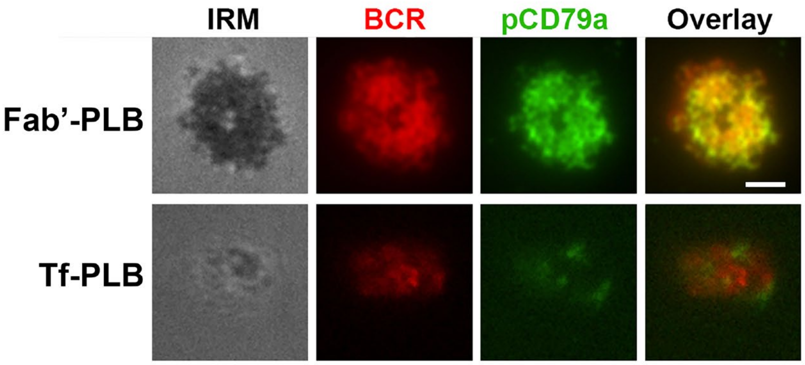


**Figure 6-figure supplement 1. Fab'-PLB, but not Tf-PLB, induces BCR clustering and phosphorylation.** WT splenic B-cells were pre-labeled with Cy3-Fab fragment of goat anti-mouse IgM+G at a concentration of 2.5 μg per $10^6$ cells at $4^{o}$C for 30 min, followed by incubation with Fab'-PLBs or Tf-PLBs for 5 min at 37°C. Cells were fixed, permeabilized, stained for pCD79a, and imaged using IRM and TIRF. Shown are representative IRM and TIRF images from three independent experiments. Scale bar, 2 μm.

### Figure 6-S2

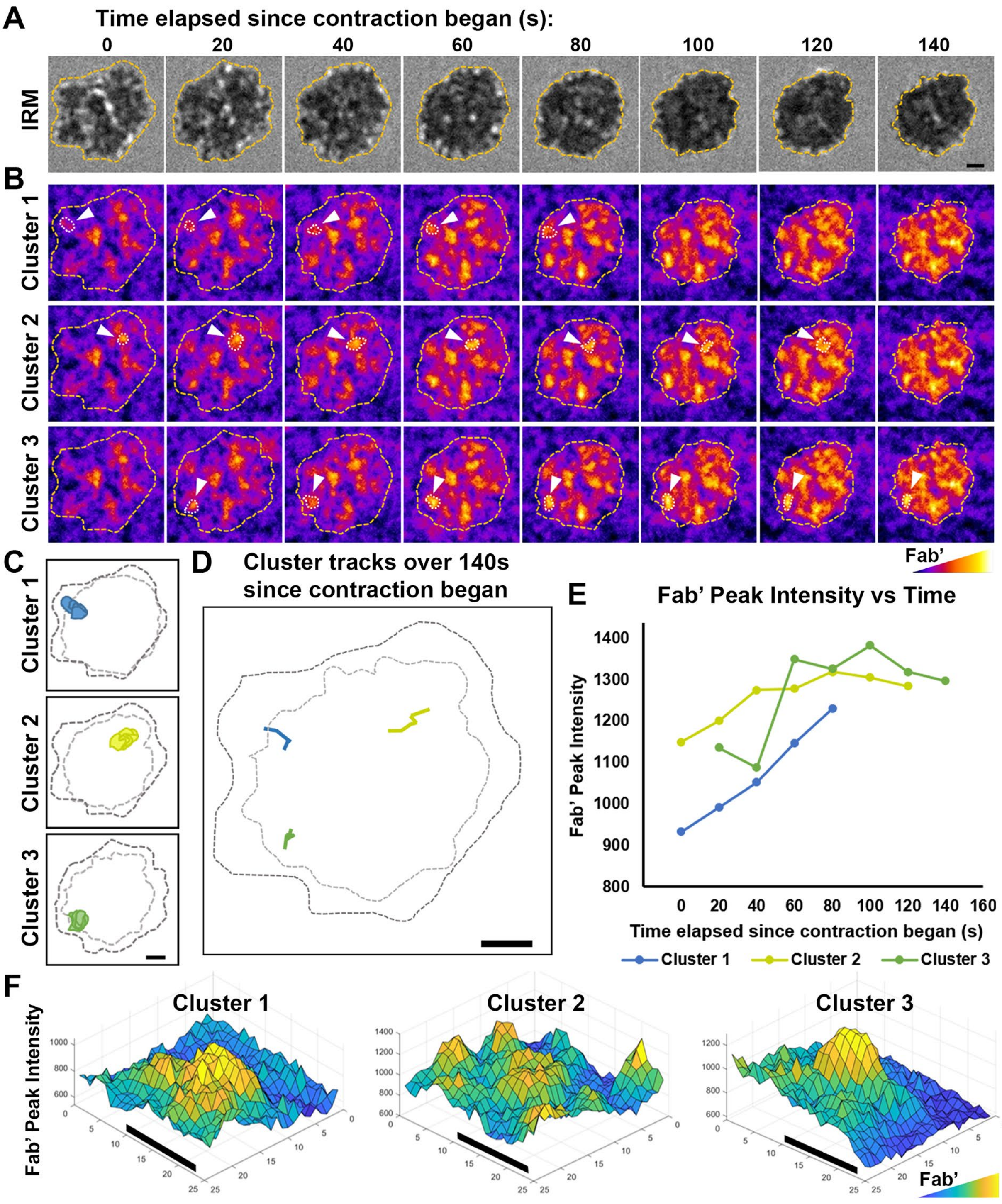


1445

**Figure 6-figure supplement 2. Tracking and analyzing AF546-Fab' clusters in the B-cell contact zone.** WT splenic B-cells were incubated with AF546-Fab'-PLB at 37°C and imaged live using IRM and TIRF. Shown are individual frames from time-lapse images of IRM (**A**) and TIRF (**B**), showing AF546-Fab' clusters within the contact zone of one DMSO-treated (vehicle control for Wisko) B-cell for 140 sec since the beginning of contraction. Fab' FI is shown as heat maps using NIH ImageJ. The boundary of the contact zone, detected using IRM images by a custom MATLAB script, is shown in yellow dashed lines. Arrows point to three representative clusters among the other clusters detected in the contact zone using custom MATLAB codes. Cluster detection masks for the three representative clusters are shown (**C**). Moving tracks for the three AF546-Fab' clusters are shown alongside the initial (black dashed lines) and final state (gray dashed lines) of the contact zone (**D**). Tracks were generated by following the peak of AF546 FI in each cluster as it moved. AF546-Fab' peak FI versus time curves for the three representative clusters are plotted over the duration that each cluster could be detected (**E**). Surface plots (2.5-D plots) of AF546-Fab' FI show a zoomed-in region consisting of each of the three AF546-Fab' clusters (**F**). Colors in (**B**) and (**F**) are scaled to AF546-Fab' FI values. Scale bars, 1 µm.
