## Supplementary material for "N-WASP-dependent branched actin polymerization attenuates B-cell receptor signaling by increasing the molecular density of receptor clusters": MATLAB scripts: analObj_v1.m

```
function [Q] = analObj_v1(RFR,bin,BG,obj_ID,FR,cFR)

cut = RFR.*bin;
Area = sum(bin(:));
[CX,CY] = Centroid(bin);
TFI = sum(cut(:));
MFI = TFI/Area;
Peak = max(cut(:));
PbM = Peak/MFI;

Q = struct('ID',obj_ID,'FR',FR,'cFR',cFR,'Area',Area,'CX',CX,'CY',CY,'TFI',TFI,'MFI',MFI,'Peak',Peak,'PbM',PbM,'BG',BG);
```
