## Supplementary material for "N-WASP-dependent branched actin polymerization attenuates B-cell receptor signaling by increasing the molecular density of receptor clusters": MATLAB scripts: MATLAB scripts_descriptions.docx

#### **Script 1:** TIRF_findCells_nd2

**To detect B cell contact zones in IRM images. This script inputs raw 16-bit .nd2 images and outputs a binary mask of all detected contact zones for further analysis.**

#### **Script 2:** TIRF_quantifyALL_nd2_3ch_v2

**To quantify all MFI and TFI data from images. This script inputs a list of binary masks of B cell contact zones, quantifies data using a sub-function (Script 2.1), and returns a table of data for each contact zone analyzed in the form of a table.**

#### **Script 2.1:** quantifyMap_nd2_3ch_v2

**To quantify contact zone area (µm2), MFI and TFI from one 3-channel 16-bit .nd2 image. This script inputs a 3-channel image consisting of one IRM channel and two fluorescent channels (labeled ‘red’ and ‘green’), as well as a mask of all B cell contact zones in the image. It calculates and outputs the contact zone area (µm2), MFI and TFI of the ‘red’ channel, and MFI and TFI of the ‘green’ channel.**

#### **Script 3:** TIRFvideo_v6

**To analyze time-lapse images and acquire Area, MFI and TFI of two fluorescent channels, in a frame-by-frame manner, for each B cell. This MATLAB liveScript is composed of several sections with some sub-functions (Scripts 3.1-3.4) It inputs several time-lapse 16-bit .tiff images consisting of 3 channels: IRM, and two fluorescent channels, each containing one B cell (cropped out). It outputs contact zone area (µm2), MFI and TFI in the form of an excel file with each sheet displaying data for multiple cells.**

#### **Script 3.1:** TIRFvid_getBinaries_v3

**To acquire a mask of a B cell contact zone from an IRM time-lapse image.**

#### **Script 3.2:** TIRFvid_saveIMstack_v1

**To save an image stack (time-lapse image) to a specified directory location**

#### **Script 3.3:** TIRFvid_quantifyExpt_v1

**To analyze fluorescent channel data in time-lapse images consisting of 3-channels: IRM and two fluorescent channels**

#### **Script 3.4:** TIRFvid_export2excel_v1

**To export time-lapse data for one B cell into a specified Excel file.**

#### **Script 3.5: TIRFvid_radialKymos_v1**

**To generate 8 kymographs from time-lapse images of a B cell consisting of 3 channels: IRM, and two fluorescent channels. The 8 kymographs are radially equally spaced. Inputs ‘Vdata’ acquired from Scripts 3.1, 3.2 and 3.3, and writes a .tiff image consisting of the 8 kymographs to the same location as the original raw data (time-lapse multi-channel image).**

#### **Script 4: TIRF_quantifyClusters_nd2_3ch_v3**

**To identify BCR clusters and signaling molecule puncta from 16-bit .nd2 images consisting of 3 channels: IRM, AF546-Fab’, and (stained) signaling molecule. Inputs B cell contact zone masks in the form of ‘MAP’ (output of Script 1). Utilizes a sub-function (Script 4.1) to quantify various parameters including Fab peak intensity, Fab MFI, and signaling molecule MFI in each detected cluster, and returns the data as a table.**

#### **Script 4.1:** quantifyClustersFOV_nd2_3ch_v9

**To identify BCR clusters and signaling molecule puncta from one 16-bit .nd2 image consisting of 3 channels: IRM, AF546-Fab’, and (stained) signaling molecule.**

#### **Script 5:** TIRFvid_vTrack_v1

**To analyze individual BCR clusters in time-lapse images. This MATLAB script is composed of several sections with some sub-functions (Scripts 5.1 and 5.2). It inputs ‘Vdata’ acquired using Script 3, identifies AF546-Fab clusters in one cropped B cell for every frame in the original 16-bit .tiff time-lapse image, generates tracks connecting the detected clusters, analyzes the peak intensity and MFI of AF546 in each cluster, and returns the output as a table.**

#### **Script 5.1:** trackObj_v1

**To track detected clusters by creating frame-by-frame association between two consecutive frames.**

#### **Script 5.2:** analObj_v1

**To analyze detected clusters and output several measured parameters including AF546 peak intensity and MFI in that cluster.**

#### **Script 6:** openND2_still

**To input multi-channel 16-bit .nd2 images (used as a sub-function in some of the above scripts)**

#### **Script 7:** openTIF_vid

**To input multi-channel 16-bit .tiff time-lapse images (used as a sub-function in some of the above scripts)**
