## Supplementary material for "N-WASP-dependent branched actin polymerization attenuates B-cell receptor signaling by increasing the molecular density of receptor clusters": MATLAB scripts: openND2_still.m

```
function [RES] = openND2_still (file_loc)

%Auto-detects number of channels in a (non-Zstack & non-video) image and
%returns as a 3-dim matrix (x,y,c).

C = bfopen_silent(file_loc);
C1 = C{1,1};
CL = size(C1,1);

RES = C1{1,1};

if CL>1
    for i = 2:1:CL
        RES(:,:,i) = C1{i,1};
    end
end

end
    
```
