## Supplementary material for "N-WASP-dependent branched actin polymerization attenuates B-cell receptor signaling by increasing the molecular density of receptor clusters": MATLAB scripts: openTIF_vid.m

```
function [RES] = openTIF_vid (file_loc,num_channels)

%Returns [1xN] cell containing 'N' 3-dim matrices (x,y,t), where 'N' is the
%number of channels.

C = bfopen_silent(file_loc);
C1 = C{1,1};
CL = size(C1,1);

if rem(CL,num_channels)>0
    error('ERROR! Check your input.');
else
    T = CL/num_channels;
    RES = cell(1,num_channels);
    samp = C1{1,1};
    for j = 1:num_channels
        RESch = zeros(size(samp,1),size(samp,2),T);
        for i = 1:T
            RESch(:,:,i) = C1{(num_channels*(i-1)+j),1};
        end
        RES{1,j} = RESch;
    end
end

end
```
