## Supplementary material for "N-WASP-dependent branched actin polymerization attenuates B-cell receptor signaling by increasing the molecular density of receptor clusters": MATLAB scripts: quantifyClustersFOV_nd2_3ch_v9.m

```
function [RES1,RES2,CRES,c_sten,n] = quantifyClustersFOV_nd2_3ch_v9(map, dir_loc, CC)

%Designed for New NIKON Microscope Images (.nd2)

%Inputs a stencil and its directory location to quantify the corresponding
%fluorescent images. Produces a struct array of data from each cell object
%in the stencil

locL = length(dir_loc);
bksl = locL;
COUNT = 1;
for i = locL:-1:1
    if dir_loc(i)=='\'
        bksl(COUNT) = i;
        COUNT = COUNT+1;
    else
        continue;
    end
end
bksl_1 = bksl(1);
bksl_2 = bksl(2);
well = dir_loc(bksl_2+1:bksl_1-1);%records the time-point or well

%disp(["Well = " well]);

imbw = map;
pix_size = 0.072; %pixel size (physical length of one pixel) in micron
%For 60X lens and 1.0X mag, it is 0.108.
%For 60X lens and 1.0X mag, it is 0.072.
pix_area = 1/(pix_size*pix_size); %pixel to area convertion ratio

c_sten = cell(2,1);
[imlb, n] = bwlabel(imbw);%label objects in final logical matrix

IP = open2Dmc_img([dir_loc, '.nd2']);
imr = IP(:,:,2);%inputs red image (BCR)
img = IP(:,:,3);%inputs green image (signaling molecule)

RES1 = struct();
RES2 = struct();
CRES = struct();%creates a struct to store results
count_1 = 1;
count_2 = 1;
cnt = 1;

D = 140;
for i = 1:n
    I = (imlb==i);%isolate each cell object
    Parea = bwarea(I);
    [cmx,cmy] = Centroid(I);
    cmx = round(cmx);
    cmy = round(cmy);
    imI = I((cmx-(D/2)) : (cmx+(D/2)-1),(cmy-(D/2)) : (cmy+(D/2)-1));
    imR = imr((cmx-(D/2)) : (cmx+(D/2)-1),(cmy-(D/2)) : (cmy+(D/2)-1));
    imR = double(imR);
    imG = img((cmx-(D/2)) : (cmx+(D/2)-1),(cmy-(D/2)) : (cmy+(D/2)-1));
    imG = double(imG);
    
    R_bg = [];
    G_cz = [];
    for j = 1:D
        for k = 1:D
            if imI(j,k)==1
                G_cz = [G_cz imG(j,k)];
            else
                R_bg = [R_bg imR(j,k)];
            end
        end
    end
    th_R = [4.1 3.9 3.7 3.5 3.3 3.1 2.9 2.7 2.5 2.3 2.1 1.9 1.7 1.5 1.3 1.1]*median(R_bg);%  <-----EDIT BCR THRESHOLDs!!!
    %th_R = [1.5 1.3]*median(R_bg);%  <-----EDIT BCR THRESHOLDs!!!
    th_G = 1.3*median(G_cz);%  <-----EDIT Green ("p-sig-mol") THRESHOLD!!!
    BCR_norm_frac = median(R_bg)/100;
    
    RMask = cell(1,size(th_R,2));
    th_R1 = th_R(1);
    imRi1 = double(imR>th_R1);
    imRi1 = imRi1.*imI;
    imRi1 = imfill(imRi1,'holes');
    imRi1 = bwareaopen(imRi1,3,4);
    se = strel('disk',1,0);

    [imlb1,n1] = bwlabel(imRi1);
    for j = 1:n1
        obj = imlb1==j;
        if sum(obj(:))<10
            imRi1 = imRi1 - obj;
            imRi1 = imRi1 + imdilate(obj,se);
        end
    end
    Mask1 = imerode(imRi1,se);
    Mask1 = imdilate(Mask1,se);
    RMask{1,1} = Mask1;
    R_Mask = Mask1*size(th_R,2);

    for k = 2:size(th_R,2)
        th_R2 = th_R(k);
        imRi2 = double(imR>th_R2);
        imRi2 = imRi2.*imI;
        imRi2 = imfill(imRi2,'holes');
        imRi2 = bwareaopen(imRi2,3,4);
        [imlb2,n1] = bwlabel(imRi2);
        for j = 1:n1
            obj = imlb2==j;
            overlap = obj.*imRi1;
            objArea = sum(obj(:));
            if sum(overlap(:)>0)||(objArea>1000)%Area upper-cutoff
                imRi2 = imRi2 - obj;
            else
                if sum(obj(:))<10
                    imRi2 = imRi2 - obj;
                    imRi2 = imRi2 + imdilate(obj,se);
                end
            end
        end
        imRi1 = imRi1 + imRi2;
        Mask2 = imerode(imRi2,se);
        Mask2 = imdilate(Mask2,se);
        RMask{1,k} = Mask2;
        R_Mask = (R_Mask + (Mask2*(size(th_R,2)+1-k)));
    end

    imGi = double(imG>th_G);
    imGi = imGi.*imI;
    imGi = imfill(imGi,'holes');
    imGi = bwareaopen(imGi,3,4);
    imGi = imerode(imGi,se);
    imGi = imdilate(imGi,se);

    GMask = imGi;
    
    RGMask = R_Mask+GMask;
    
    if (sum(RGMask(:))>0)
        for j = 1:size(th_R,2)
            Mask_i = RMask{1,j};
            [imRlb,n2] = bwlabel(Mask_i);%checking clusters...
            for k = 1:n2
                Ci = double(imRlb==k);
                Cr = imR.*Ci;
                Cg = imG.*Ci;
                Area = sum(Ci(:));
                TFI = sum(Cr(:))/BCR_norm_frac;
                MFI = TFI/Area;
                Peak = max(Cr(:))/BCR_norm_frac;
                PbM = Peak/MFI;
                pMFI = sum(Cg(:))/Area;
                pMFIR = pMFI/MFI;

                RES1(count_1).Well = well;%stores well (time-point) info
                RES1(count_1).Cell = CC-1+i;%stores cell number
                RES1(count_1).Type = j;%stores BCR cluster type: 1 = High; 2 = Med; 3 = Low...
                RES1(count_1).Area = Area;%stores BCR cluster size in pix
                RES1(count_1).TFI = TFI;
                RES1(count_1).MFI = MFI;
                RES1(count_1).Peak = Peak;
                RES1(count_1).PbM = PbM;
                RES1(count_1).pMFI = pMFI;
                RES1(count_1).pMFIR = pMFIR;
                count_1 = count_1+1;
            end
        end
        
        [imGlb,n2] = bwlabel(GMask);%checking p-sig-mol puncta...
        for k = 1:n2
            Ci = double(imGlb==k);
            Cr = imR.*Ci;
            Cg = imG.*Ci;
            Area = sum(Ci(:));
            TFI = sum(Cg(:));
            MFI = TFI/Area;
            Peak = max(Cg(:));
            PbM = Peak/MFI;
            rPeak = max(Cr(:))/BCR_norm_frac;
            rTFI = sum(Cr(:))/BCR_norm_frac;
            rMFI = sum(Cr(:))/(Area*BCR_norm_frac);
            rPbM = rPeak/rMFI;
            pMFIR = MFI/rMFI;

            RES2(count_2).Well = well;%stores well (time-point) info
            RES2(count_2).Cell = CC-1+i;%stores cell number
            RES2(count_2).Type = 0;%stores BCR cluster type: 0 = p-sig-mol...
            RES2(count_2).Area = Area;%stores puncta size in pix
            RES2(count_2).TFI = rTFI;
            RES2(count_2).MFI = rMFI;
            RES2(count_2).Peak = rPeak;
            RES2(count_2).PbM = rPbM;
            RES2(count_2).pTFI = TFI;
            RES2(count_2).pMFI = MFI;
            RES2(count_2).pPeak = Peak;
            RES2(count_2).pPbM = PbM;
            RES2(count_2).pMFIR = pMFIR;
            
            count_2 = count_2+1;
        end
        
        c_sten{1,i} = R_Mask;
        c_sten{2,i} = GMask;
        
        CRES(cnt).Well = well;%stores well (time-point) info
        CRES(cnt).Cell = CC-1+i;%stores cell number
        CRES(cnt).Area = (Parea/pix_area);%stores IRM area
        cnt = cnt+1;
    else
        c_sten{1,i} = 0*imRi1;
        c_sten{2,i} = 0*imRi1;
    end
end

end
```
