## Supplementary material for "N-WASP-dependent branched actin polymerization attenuates B-cell receptor signaling by increasing the molecular density of receptor clusters": MATLAB scripts: quantifyMap_nd2_3ch_v2.m

```
function [RES] = quantifyMap_nd2_3ch_v2(M, dir_loc)

%Designed for New NIKON Microscope Images (.nd2)

%Inputs a stencil and its directory location to quantify the corresponding
%fluorescent images. Produces a struct array of data from each cell object
%in the stencil

locL = length(dir_loc);
bksl = locL;
count = 1;
for i = locL:-1:1
    if dir_loc(i)=='\'
        bksl(count) = i;
        count = count+1;
    else
        continue;
    end
end
bksl_1 = bksl(1);
bksl_2 = bksl(2);
well = dir_loc(bksl_2+1:bksl_1-1);%records the time-point or well

imbw1 = M;
pix_size = 0.072; %pixel size (physical length of one pixel) in micron
%For 60X lens and 1.0X mag, it is 0.108.
%For 60X lens and 1.0X mag, it is 0.072.
%For 100X lens and 1.0X mag, it is 0.065.
pix_area = 1/(pix_size*pix_size); %pixel to area convertion ratio

[imlb, n] = bwlabel(imbw1);%label objects in final logical matrix

IP = open2Dmc_img([dir_loc, '.nd2']);
%imp0 = IP(:,:,2);%inputs purple image (channel 1)
imr0 = IP(:,:,2);%inputs red image (channel 1)
img0 = IP(:,:,3);%inputs green image (channel 2)

%imgbg = 65535*double(adaptthresh(img0,0.65));
%imrbg = 65535*double(adaptthresh(imr0,0.65));
%impbg = 65535*double(adaptthresh(imp0,0.65));%thresholds

img = double(img0);
imr = double(imr0);
%imp = double(imp0);%converts image:must be done after adaptthresh:

%img = img-imgbg;
%imr = imr-imrbg;%negates threshold

RES = struct();%creates a struct to store results

for i = 1:n
    I = (imlb==i);%isolate each cell object
    
    se1 = strel('disk',10,0);
    se2 = strel('disk',100,0);
    ibin_ex1 = imdilate(I,se1);
    ibin_ex2 = imdilate(I,se2);
    ibin_bg = ibin_ex2 - ibin_ex1;
    I_bg = ibin_bg-(ibin_bg.*(imdilate(imbw1,se1)));
    
    imgi = img.*I;
    imri = imr.*I;
    %impi = imp.*I;
    imgibg = img.*I_bg;
    imribg = imr.*I_bg;
    %impibg = imp.*I_bg;
    Parea = bwarea(I);
    Pbg = bwarea(I_bg);
    
    %pMean = sum(impi(:))/Parea - sum(impibg(:))/Pbg;
    %pTotal = pMean*Parea;
    
    gMean = sum(imgi(:))/Parea - sum(imgibg(:))/Pbg;
    gTotal = gMean*Parea;
    
    rMean = sum(imri(:))/Parea - sum(imribg(:))/Pbg;
    rFold = (sum(imri(:))/Parea)/(sum(imribg(:))/Pbg);
    rTotal = rMean*Parea;
    
    RES(i).Well = well;%stores well (time-point) info
    
    RES(i).Area = (Parea/pix_area);%stores IRM area
    
    %if pMean < 0
    %    RES(i).P_TFI= 0;
    %    RES(i).P_MFI = 0;
    %else
    %    RES(i).P_TFI=pTotal;%stores TFI of Purple
    %    RES(i).P_MFI=pMean;%stores MFI of Purple
    %end
    RES(i).P_TFI = 0;
    RES(i).P_MFI = 0;
    
    if gMean < 0
        RES(i).G_TFI= 0;
        RES(i).G_MFI = 0;
    else
        RES(i).G_TFI=gTotal;%stores TFI of Green
        RES(i).G_MFI=gMean;%stores MFI of Green
    end
    
    if rMean < 0
        RES(i).R_TFI= 0;
        RES(i).R_MFI = 0;
    else
        RES(i).R_TFI=rFold;%stores Fold increase in Red MFI
        RES(i).R_MFI=rMean;%stores MFI of Red
    end
    
    if (rMean < 0)||(gMean < 0)
        RES(i).PCC = 0;
    else
        RES(i).PCC = corr2D_PCC(I,(img-sum(imgibg(:))/Pbg),(imr-sum(imribg(:))/Pbg));%stores PCC of purple & green...
    end
    
end%quantified individual cell objects and stored values in the struct

end
```
