## Supplementary material for "N-WASP-dependent branched actin polymerization attenuates B-cell receptor signaling by increasing the molecular density of receptor clusters": MATLAB scripts: TIRF_findCells_nd2.m

```
%Designed for New NIKON Microscope Images (.nd2)

%Inputs a struct of masks (MAP) and directory location of an IRM image.
%Stores the newly obtained logical (binary) mask, well ID and the image name in the struct MAP and returns it.

%Circles objects and asks user to remove if necessary. Also has two different modes: "z" and "x".
%"z" enables user to select objects instead of removing them. The code removes all other unselected objects (reversemode).
%"x" enables user to manually draw objects.

function [MAP] = TIRF_findCells_nd2(MAP,dir_loc)

locL = length(dir_loc);
bksl = locL;
count = 1;
for i = locL:-1:1
    if dir_loc(i)=='\'
        bksl(count) = i;
        count = count+1;
    else
        continue;
    end
end
bksl_1 = bksl(1);
bksl_2 = bksl(2);
well = dir_loc(bksl_2+1:bksl_1-1);%records the time-point or well

disp(' ');
disp('*Looking for objects...')

IP = openND2_still([dir_loc, '.nd2']);
im = IP(:,:,1);%inputs IRM image
im = imadjust(im);
im1 = (imadjust(uint16(bpass(65535-im,1,150,20))));%inverts and band-passes IRM image
t = 40000;%set threshold value
imbw = im1>t;%apply threshold
se = strel('disk',1,0);
se2 = strel('disk',2,0);
imbw1 = bwareaopen((imerode((imdilate(imbw,se2)),se)),750);%clean up noise
Rd = 1000;%drop-off radius
[imlb, n] = bwlabel(imbw1);%label objects in logical matrix
for k = 1:1:n
    I = (imlb==k);%isolate each cell object
    if beyond_dropoff_v1(I,Rd) == 1%check if the centroid of the cell is too far
        imbw1 = (imbw1 - I);%omit cell if too far
    else
        continue;
    end
end%excludes cells beyond the drop-off radius
imbw1 = imfill(imbw1,'holes');%fills holes

f = 1;%set flag
while f==1

    [imlb, n] = bwlabel(imbw1);%label objects in logical matrix
    B = cell(1,n);

    subplot(1,2,1);
    h1 = imshow(imadjust(im));

    subplot(1,2,2);
    imshow(imadjust(im));
    hold on;

    for i = 1:n
        I = (imlb==i);%isolate each cell object
        Bound = bwboundaries(I);
        B(1,i) = Bound(1,1);%get boundaries of each cell
        l = size(B{1,i});

        for j = 1:l
            plot(B{1,i}(:,2), B{1,i}(:,1), 'r.', 'MarkerSize', 2);
        end%plots boundaries
        [cm_x, cm_y] = Centroid(I);
        text((cm_y-20),cm_x,num2str(i),'Color','green','FontSize',10);%labels cells on image
    end

    x = 1024;
    y = 1024;
    ang=0:0.01:2*pi;
    xp=Rd*cos(ang);
    yp=Rd*sin(ang);
    plot(x+xp,y+yp,'m.');%plots 'drop-off circle'
    hold off;

    %now check if user is happy with labelled objects...
    InfoBox = uicontrol('style','text');
    set(InfoBox,'String','To remove objects, enter their corresponding numbers separated by spaces. If everything looks good, enter 0. ');
    set(InfoBox,'Position',[20,20,700,50]);%puts a sentence in the figure with instructions for the user

    disp('To remove objects, enter their corresponding numbers separated by spaces.');
    disp('You can also press "z" or "x" to enter one of two add-on modes.');
    disp('"z" = Keep selected objects (reverse mode). "x" = Draw cells manually.');
    prompt = 'If everything looks good, enter 0. ';%also prompts user for input(s)
    omit = input(prompt,'s');
    if (strcmp(omit,'0')~=1)
        if ((strcmp(omit,'x')~=1)&&(strcmp(omit,'z')~=1))%For default mode: "Remove selected."
            omit = ([omit ' ']);
            omit_num = 1;
            omit_val = ['0'];
            omit_List = [];
            omit_str_len = length(omit);
            for i = 1:1:omit_str_len%acquires list from string input
                if (omit(i) ~= ' ')
                    omit_val = ([omit_val omit(i)]);
                elseif (omit(i) == ' ')
                    omit_List(omit_num) = str2num(omit_val);
                    omit_num = omit_num + 1;
                    omit_val = ['0'];
                end
            end

            for i = 1:1:size(omit_List,2)%omits objects
                omit_obj = omit_List(i);
                if (omit_obj == 0)
                    continue;
                elseif ((omit_obj > n)||(omit_obj <=0))
                    error(['ERROR! Could not find object number ' num2str(omit_obj)]);%throws error for out-of-bound input
                end
                imbw1 = imbw1 - bwselect(imbw1,B{1,omit_obj}(1,2),B{1,omit_obj}(1,1));%remove unwanted object and loop...
                disp([num2str(omit_obj) ' removed.']);
            end

            if omit_List(size(omit_List,2))==0
                f = 0;
            end

        elseif (strcmp(omit,'z')==1)%For 'z' mode: "Keep objects, remove all else."

            prompt = 'Enter the objects you want to keep, separated by spaces. ';
            omit = input(prompt,'s');
            omit = ([omit ' ']);
            omit_num = 1;
            omit_val = ['0'];
            omit_List = [];
            omit_str_len = length(omit);
            for i = 1:1:omit_str_len%acquires list from string input
                if (omit(i) ~= ' ')
                    omit_val = ([omit_val omit(i)]);
                elseif (omit(i) == ' ')
                    omit_List(omit_num) = str2num(omit_val);
                    omit_num = omit_num + 1;
                    omit_val = ['0'];
                end
            end
            imbw2 = logical(zeros(size(imbw1,1),size(imbw1,2)));
            for i = 1:1:size(omit_List,2)%selects objects
                omit_obj = omit_List(i);
                if (omit_obj == 0)
                    continue;
                elseif ((omit_obj > n)||(omit_obj <=0))
                    error(['ERROR! Could not find object number ' num2str(omit_obj)]);%throws error for out-of-bound input
                end
                imbw2 = imbw2 + bwselect(imbw1,B{1,omit_obj}(1,2),B{1,omit_obj}(1,1));%add object to new logical and loop...
                disp([num2str(omit_obj) ' saved.']);
            end
            imbw1 = imbw2;
            if omit_List(size(omit_List,2))==0
                f = 0;
            end

        elseif (strcmp(omit,'x')==1)%For 'x' mode: "Draw objects manually."
            add_Fg = 1;
            add_count = 1;
            add_mask = zeros(size(im,1),size(im,2));
            add_IM = double(im);
            add_sillh = imbw1.*(max(add_IM));
            add_IM = (add_IM.*(1-imbw1))+add_sillh;
            imshow(imadjust(uint16(add_IM)));
            hold on;

            while(add_Fg == 1)
                Mobj = imfreehand;
                new_add = createMask(Mobj,h1);
                add_mask(:,:,add_count) = new_add;
                add_count = add_count + 1;
                prompt = 'If you want to circle more cells, enter "x". If not, enter anything else.';
                rep = input(prompt,'s');
                if strcmp(rep,'x')~=1
                    add_Fg = 0;
                end
            end
            hold off;

            add_mask_all = sum(add_mask,3);
            check_ovrlp = (imbw1.*add_mask_all);
            if (sum(check_ovrlp(:))>0)
                error('ERROR! Some of your circled cells overlap with existing objects on this image...');
            end
            imbw1 = imbw1 + add_mask_all;
            disp(['Additional ' num2str(add_count-1) ' objects were added to the mask.']);
        end
        
    elseif size(omit) == 0
        disp('You did not enter any text...');%throws error for blank input

    elseif strcmp(omit,'0')
        f = 0;%set flag to exit while loop and save stencil/mask
        break;
    end
end
    
M = logical(imbw1);

[imlb2, nf] = bwlabel(M);

ST = struct('Well',well,'fname',dir_loc,'stencil',M);%temporary data element to be stored in a struct

flag = 0;%checks for repetition before saving data
sz = size(MAP,2);
for i = sz:-1:2
    if strcmp(MAP(i).fname,dir_loc)==1
        flag = 1;
        MAP(i) = ST;
        disp('Mask overwritten!');
        break;
    end
end
if flag==0
    MAP = ([MAP,ST]);
end%stores well, filename and stencil in the MAP struct

disp('*****COMPLETED!!!');
disp(['Number of objects saved = ' num2str(nf)]);

end
```
