## Supplementary material for "N-WASP-dependent branched actin polymerization attenuates B-cell receptor signaling by increasing the molecular density of receptor clusters": MATLAB scripts: TIRF_quantifyALL_nd2_3ch_v2.m

```
function [Q1] = TIRF_quantifyALL_nd2_3ch_v2(MAP)

%Designed for New NIKON Microscope Images (.nd2)

%Inputs a struct MAP of stencils and the directory location of the data
%folder for a whole TIRF experiment. Accesses fluorescent images for each
%stencil, quantifies data, removes outliers and stores it in an output
%struct

Q1 = struct([]);
Q = MAP;

WL = {' '};
count = 1;
for i = 2:length(Q)
    flag = 0;
    for j = 1:size(WL,2)
        if strcmp(WL{1,j},Q(i).Well)
            flag = 1;
        end
    end
    if flag == 0
        WL{1,count} = Q(i).Well;
        count = count+1;
    end
end
%generates a list of 'wells' from the struct input

for i = 1:size(WL,2)
    well = WL{1,i};
    ST = getWells(Q,well);%gets all stencils from one well
    
    W = struct([]);
    for j = 1:size(ST,2)
        dir_loc = ST(j).fname;%gets directory location from input struct
        M = ST(j).stencil;%gets stencil from input struct
        if max(M(:))==0
            continue;
        else
            R = quantifyMap_nd2_3ch_v2(M,dir_loc);%computes and stores values as a Struct R
            W = [W, R];%appends data from one set of images to the data for the whole timepoint (well)
        end
    end
    %W = processData_v3(W);
    Q1 = [Q1,W];%appends data from one well to the main data Struct (output)
end

viewStruct(Q1);

end
```
