## Supplementary material for "N-WASP-dependent branched actin polymerization attenuates B-cell receptor signaling by increasing the molecular density of receptor clusters": MATLAB scripts: TIRF_quantifyClusters_nd2_3ch_v3.m

```
function [Q1,Q2,Qcells,cMAP] = TIRF_quantifyClusters_nd2_3ch_v3(MAP)

%Designed for New NIKON Microscope Images (.nd2)

%Inputs a struct MAP of stencils and the directory location of the data
%folder for a whole TIRF experiment. Accesses fluorescent images for each
%stencil, quantifies data, removes outliers and stores it in an output
%struct

Q1 = struct([]);
Q2 = struct([]);
Qcells = struct([]);
M = MAP;

cMAP = MAP(1);

WL = {' '};
count = 1;
for i = 2:length(M)
    flag = 0;
    for j = 1:size(WL,2)
        if strcmp(WL{1,j},M(i).Well)
            flag = 1;
        end
    end
    if flag == 0
        WL{1,count} = M(i).Well;
        count = count+1;
    end
end
%generates a list of 'wells' from the struct input

for i = 1:size(WL,2)
    well = WL{1,i};
    ST = getWells(M,well);%gets all stencils from one well
    cm = ST;
    W1 = struct([]);
    W2 = struct([]);
    CW = struct([]);
    Ccount = 1;
    for j = 1:size(ST,2)
        dir_loc = ST(j).fname;%gets directory location from input struct
        map = ST(j).stencil;%gets stencil from input struct
        if max(map(:))==0
            continue;
        else
            [R1,R2,CR,c_sten,CcountI] = quantifyClustersFOV_nd2_3ch_v9(map,dir_loc,Ccount);%computes and stores values as a Struct R
            if (~isempty(fieldnames(R1)))||(~isempty(fieldnames(R2)))
                if (~isempty(fieldnames(R1)))
                W1 = [W1, R1];%appends data from one set of images to the data for the whole timepoint (well)
                end
                if (~isempty(fieldnames(R2)))
                    W2 = [W2, R2];%appends data from one set of images to the data for the whole timepoint (well)
                end
                if (~isempty(fieldnames(CR)))
                    CW = [CW, CR];
                end
                Ccount = Ccount + CcountI;
            end
            cm(j).stencil = c_sten;
        end
    end
    %W = processData_v3(W);
    if (~isempty(fieldnames(W1)))
        Q1 = [Q1,W1];%appends data from one well to the main data Struct (output)
    end
    if (~isempty(fieldnames(W2)))
        Q2 = [Q2,W2];%appends data from one well to the main data Struct (output)
    end
    if (~isempty(fieldnames(CW)))
        Qcells = [Qcells,CW];%appends data from one well to the main data Struct (output)
    end
    cMAP = [cMAP,cm];
end

end
```
