## Supplementary material for "N-WASP-dependent branched actin polymerization attenuates B-cell receptor signaling by increasing the molecular density of receptor clusters": MATLAB scripts: TIRFvid_export2excel_v1.m

```
function [OP] = TIRFvid_export2excel_v1(Qdata,out_excel,out_cell)

Vmeta = Qdata(1).Mask_IRM;
frameRATE = Vmeta.frameRATE;
FN = fieldnames(Qdata);
div = 0;
for i = 1:size(FN,1)
    if (strcmp(FN{i,1},'Area')||strcmp(FN{i,1},'MFI_out'))
        div = i;
        break;
    end
end
OP = FN(div:(length(FN)-1),1);

cell_prop = isstrprop(out_cell,'alpha');
cell_X = out_cell(1:sum(cell_prop));
cell_Y = out_cell(sum(cell_prop)+1:length(out_cell));
if length(cell_X)==1
    cXn = double(cell_X)-64+1;
else
    cXn = ((double(cell_X)-64)*[26; 1])+1;
end
Y_loc = num2str(str2num(cell_Y)+1);
max_fr = 0;

for k = 2:size(Qdata,2)
    cellID = {Qdata(k).cellID};
    Mask_IRM = Qdata(k).Mask_IRM;
    len_fr = size(Mask_IRM,3);
    if len_fr>max_fr
        max_fr = len_fr;
    end
    active_sheet = 1;
    for j = div:size(FN,1)
        Djk = getfield(Qdata(k),FN{j,1});
        if cXn<=26
            X_loc = char(cXn+64);
        else
            X_loc = [char(floor(cXn/26)+64) char(rem(cXn,26)+64)];
        end
        cell = [X_loc cell_Y];
        record_excel(cellID,out_excel,active_sheet,cell);
        cell = [X_loc Y_loc];
        record_excel(Djk',out_excel,active_sheet,cell);
        active_sheet = active_sheet + 1;%next sheet/field
    end
    cXn = cXn + 1;%next crop/shift right
end

Time = [1:1:max_fr];
Time = Time/frameRATE;
Time = [0 Time];
if length(cell_X)==1
    cXn = double(cell_X)-64;
else
    cXn = ((double(cell_X)-64)*[26; 1]);
end
active_sheet = 1;
for j = div:(size(FN,1)-1)
    if cXn<=26
        X_loc = char(cXn+64);
    else
        X_loc = [char(floor(cXn/26)+64) char(rem(cXn,26)+64)];
    end
    cell = [X_loc cell_Y];
    record_excel({['Time(s)']},out_excel,active_sheet,cell);
    cell = [X_loc Y_loc];
    record_excel(Time',out_excel,active_sheet,cell);
    active_sheet = active_sheet + 1;%next sheet/field
end
cell = [X_loc Y_loc];
key_other = {'Start lag(s)'};
record_excel(key_other,out_excel,active_sheet,cell);

end
```
