## Supplementary material for "N-WASP-dependent branched actin polymerization attenuates B-cell receptor signaling by increasing the molecular density of receptor clusters": MATLAB scripts: TIRFvid_getBinaries_v3.m

```
function [M3] = TIRFvid_getBinaries_v3(filename,num_channels,actin_channel,IRM_thresh,Actin_thresh,IRM_sizeCUToff,Actin_sizeCUToff)

IP = openTIF_vid(filename,num_channels);
Vi = IP{1,1};
Vf = IP{1,actin_channel};
Vi = uint16(Vi);
Vf = uint16(Vf);
tn = size(Vf,3);
manual_thresh = Actin_thresh;

%Detect footprints in last frame:

FRi = Vi(:,:,tn);% (IRM)
FRf = Vf(:,:,tn);% (actin)
THi = IRM_thresh;
%THf = Actin_thresh;
ISCO = IRM_sizeCUToff;
ASCO = Actin_sizeCUToff;

imbw1 = FRi<THi;%apply threshold (IRM)
se = strel('disk',3,0);%for smoothing
se3 = strel('disk',3,0);%for dilation of prev_fr
imbw1 = imerode((imdilate(imbw1,se)),se);%dilation/erosion=smoothing (IRM)
imbw1 = bwareaopen(imbw1,ISCO);%cleans noise (IRM)
imbw1 = imfill(imbw1,'holes');%fills holes (IRM)
prev_fr1 = imdilate(imbw1,se3);

[counts,x] = imhist(uint16(FRf),65535);
THf = otsuthresh(counts)*65535;

if manual_thresh>0
    THf = manual_thresh;
end

imbw2 = FRf>THf;%apply threshold (actin)
imbw2 = imerode((imdilate(imbw2,se)),se);%dilation/erosion=smoothing (actin)
imbw2 = bwareaopen(imbw2,ASCO);%cleans noise (actin)
imbw2 = imfill(imbw2,'holes');%fills holes (actin)
prev_fr2 = imbw2;

imbw3 = (imbw1+imbw2)>0;
imbw3 = imerode((imdilate(imbw3,se)),se);
imbw3 = imfill(imbw3,'holes');%fills holes

%[imlb,N] = bwlabel(imbw3);
%imbw3 = imbw3 - (imlb==2);

% Detect footprints in remaining frames until the first:
M1 = zeros(size(Vf));
M2 = M1;
M3 = M1;
M1(:,:,tn) = imbw1;
M2(:,:,tn) = imbw2;
M3(:,:,tn) = imbw3;
for i = tn-1:-1:1
    FRi = Vi(:,:,i);
%    THi = 65535*graythresh(FRi);%set threshold (IRM)
    imbw1 = FRi<THi;%apply threshold (IRM)
    imbw1 = imerode((imdilate(imbw1,se)),se);%dilation/erosion=smoothing (IRM)
    imbw1 = bwareaopen(imbw1,ISCO);%cleans noise (IRM)
    imbw1 = imfill(imbw1,'holes');%fills holes (IRM)
    [imlb1,n] = bwlabel(imbw1);
    if n>1
        for j = 1:n
            obj = imlb1==j;
            [cm_x, cm_y] = Centroid(obj);
            if prev_fr1(round(cm_x),round(cm_y))==0
                imbw1 = imbw1 - obj;
            end
        end
    end
    M1(:,:,i) = imbw1;
    prev_fr1 = imdilate(imbw1,se3);
    
    FRf = Vf(:,:,i);
    [counts,x] = imhist(uint16(FRf),65535);
    THf = otsuthresh(counts)*65535;
    
    if manual_thresh>0
        THf = manual_thresh;
    end

    imbw2 = FRf>THf;%apply threshold (actin)
    imbw2 = imerode((imdilate(imbw2,se)),se);%dilation/erosion=smoothing (actin)
    imbw2 = bwareaopen(imbw2,ASCO);%cleans noise (actin)
    imbw2 = imfill(imbw2,'holes');%fills holes (actin)
    [imlb2,n] = bwlabel(imbw2);
    if n>1
        for j = 1:n
            obj = imlb2==j;
            [cm_x, cm_y] = Centroid(obj);
            if prev_fr2(round(cm_x),round(cm_y))==0
                imbw2 = imbw2 - obj;
            end
        end
    end
    M2(:,:,i) = imbw2;
    prev_fr2 = imbw2;
    
    imbw3 = (imbw1+imbw2)>0;
    imbw3 = imerode((imdilate(imbw3,se)),se);
    imbw3 = imfill(imbw3,'holes');%fills holes
    
    %if i>80
    %    [imlb,N] = bwlabel(imbw3);
    %    imbw3 = imbw3 - (imlb==2);
    %end
    
    M3(:,:,i) = imbw3;

end

end
```
