## Supplementary material for "N-WASP-dependent branched actin polymerization attenuates B-cell receptor signaling by increasing the molecular density of receptor clusters": MATLAB scripts: TIRFvid_quantifyExpt_v1.m

```
function [Qdata] = TIRFvid_quantifyExpt_v1(Vdata)

Qdata = Vdata;

Vmeta = Vdata(1).Mask_IRM;
frameRATE = Vmeta.frameRATE;
pixLEN = Vmeta.pixLEN;
num_channels = Vmeta.num_channels;
actin_channel = Vmeta.actin_channel;
channels = Vmeta.channels;

for k = 2:size(Vdata,2)
    disp(['Quantifying cell ' num2str(k-1) ' of ' num2str(size(Vdata,2)-1) '...']);
    filename = Vdata(k).fName;
    IP1 = openTIF_vid(filename,num_channels);
    V_IRM = IP1{1,1};
    V_FLU1 = IP1{1,2};
    if length(channels)>2
        V_FLU2 = IP1{1,3};
    end
    tn = size(V_IRM,3);
    Mask_IRM = Vdata(k).Mask_IRM;
    Mask_act = Vdata(k).Mask_act;
    CFR = Vdata(k).CFR;
    SFR = Vdata(k).SFR;
    Q_sl = Vdata(k);
    
    % Calculate Area data:
    Area = zeros(1,size(Mask_IRM,3));
    for i = 1:tn
        bin = Mask_IRM(:,:,i);
        Area(i) = sum(bin(:));
    end
    Area = Area*(pixLEN*pixLEN/1000000);%correct pixel area to area in um squared
    Area = Area(SFR:length(Area));
    Area = [0 Area];
    if(k==2)
        Qdata(1).Area = 'Area';
    end
    Q_sl.Area = Area;%records Area

    VcI = ones(size(V_IRM));
    VcI = VcI - double(V_IRM);
    TFI1 = zeros(1,size(Mask_IRM,3));
    MFI1 = zeros(1,size(Mask_IRM,3));
    for i = SFR:tn
        bin = Mask_IRM(:,:,i);
        bin_prod = bin.*VcI(:,:,i);
        tfi1 = sum(bin_prod(:));
        TFI1(i) = tfi1;
        mfi1 = tfi1/sum(bin(:));
        MFI1(i) = mfi1;
    end
    TFIbg1 = zeros(1,size(Mask_IRM,3));
    MFIbg1 = zeros(1,size(Mask_IRM,3));
    for i = SFR:tn
        bin = Mask_IRM(:,:,i);
        binBG = 1-bin;
        bin_prod_bg = binBG.*VcI(:,:,i);
        tfibg1 = sum(bin_prod_bg(:));
        TFIbg1(i) = tfibg1;
        mfibg1 = tfibg1/sum(binBG(:));
        MFIbg1(i) = mfibg1;
    end
    MFI1_corr = MFI1 - MFIbg1;%Background correction of MFI
    MFI1_corr = MFI1_corr(SFR:length(MFI1_corr));
    MFI1_corr = [0 MFI1_corr];
    if(k==2)
        Qdata(1).IRM_MFI = 'IRM_MFI';
    end
    Q_sl.IRM_MFI = MFI1_corr;%records IRM MFI (density)

    dAdt = diff(Area);
    dAdt(1) = 0;
    dAdt = [0 dAdt];
    dAdt = dAdt*frameRATE;
    if(k==2)
        Qdata(1).dAdt = 'dAdt';
    end
    Q_sl.dAdt = dAdt;%records dA/dt
    
    %Calculate data from other fluorescent channel(s)
    for j = 2:length(channels)
        Mask = Mask_IRM;
        ch_N = channels(j).name;
        TFI1 = zeros(1,size(Mask,3));
        MFI1 = zeros(1,size(Mask,3));
        if j==2
            V_FLU = V_FLU1;
        elseif j==3
            V_FLU = V_FLU2;
        end
        for i = SFR:tn
            bin = Mask(:,:,i);
            bin_prod = bin.*V_FLU(:,:,i);
            tfi1 = sum(bin_prod(:));
            TFI1(i) = tfi1;
            mfi1 = tfi1/sum(bin(:));
            MFI1(i) = mfi1;
        end
        TFIbg1 = zeros(1,size(Mask,3));
        MFIbg1 = zeros(1,size(Mask,3));
        TFIbg1_total = TFIbg1;
        for i = SFR:tn
            bin = Mask(:,:,i);
            binBG = 1-bin;
            bin_prod_bg = binBG.*V_FLU(:,:,i);
            tfibg1 = sum(bin_prod_bg(:));
            TFIbg1(i) = tfibg1;
            mfibg1 = tfibg1/sum(binBG(:));
            MFIbg1(i) = mfibg1;
            TFIbg1_total(i) = mfibg1*(sum(bin(:)));
        end
        MFI1_corr = MFI1 - MFIbg1;%Background correction of MFI
        MFI1_corr = MFI1_corr(SFR:length(MFI1_corr));
        MFI1_corr = [0 MFI1_corr];
        if(k==2)
            if j==2
                Qdata(1).C2_MFI = [ch_N '_MFI'];
            elseif j==3
                Qdata(1).C3_MFI = [ch_N '_MFI'];
            end
        end
        if j==2
            Q_sl.C2_MFI = MFI1_corr;
        elseif j==3
            Q_sl.C3_MFI = MFI1_corr;
        end%records MFI

        TFI1_corr = TFI1 - TFIbg1_total;
        TFI1_corr = TFI1_corr(SFR:length(TFI1_corr));
        TFI1_corr = [0 TFI1_corr];
        ch_N = channels(j).name;
        if(k==2)
            if j==2
                Qdata(1).C2_TFI = [ch_N '_TFI'];
            elseif j==3
                Qdata(1).C3_TFI = [ch_N '_TFI'];
            end
        end
        if j==2
            Q_sl.C2_TFI = TFI1_corr;
        elseif j==3
            Q_sl.C3_TFI = TFI1_corr;
        end%records TFI

        TFI1 = [0 TFI1];
        TFI1 = TFI1(SFR:length(TFI1));
        dXdt = diff(TFI1);
        dXdt(1) = 0;
        dXdt = [0 dXdt];
        dXdt = dXdt*frameRATE;
        if(k==2)
            if j==2
                Qdata(1).C2_dXdt = [ch_N '_dXdt'];
            elseif j==3
                Qdata(1).C3_dXdt = [ch_N '_dXdt'];
            end
        end
        if j==2
            Q_sl.C2_dXdt = dXdt;
        elseif j==3
            Q_sl.C3_dXdt = dXdt;
        end%records dX/dt
    end
    S_lag = (SFR-CFR)/frameRATE;
    Qdata(1).S_lag = 'S_lag';
    Q_sl.S_lag = S_lag;%records S_lag
    Qdata(k) = Q_sl;
end

end
```
