## Supplementary material for "N-WASP-dependent branched actin polymerization attenuates B-cell receptor signaling by increasing the molecular density of receptor clusters": MATLAB scripts: TIRFvid_radialKymos_v1.m

```
%Load Vdata first!

for mapN = 7
    
    manual_R = 40;%Enter -1 for auto radius...
    
    filename = Vdata(mapN).fName;
    savename = [filename(1:length(filename)-4) '_kymos.tif'];

    IP = openTIF_vid(filename,3);
    T = [Vdata(mapN).SFR (Vdata(mapN).SFR-1+90)];
    Mask = Vdata(mapN).Mask_IRM;

    CMX = [];
    CMY = [];
    B = [];
    for t = T(1):1:T(2)
        Mask_fr = Mask(:,:,t);
        [cmx,cmy] = Centroid(Mask_fr);
        CMX = [CMX cmx];
        CMY = [CMY cmy];
        Bound = bwboundaries(Mask_fr);
        bound = Bound{1,1};
        Bdist = [];
        for i = 1:1:size(bound,2)
            bdist = sqrt(((bound(i,1)-cmx)*(bound(i,1)-cmx))+((bound(i,2)-cmy)*(bound(i,2)-cmy)));
            Bdist = [Bdist bdist];
        end
        B = [B max(Bdist)];
    end
    cmx = round(mean(CMX));
    cmy = round(mean(CMY));
    if manual_R>0
        R = manual_R;
    else
        R = 5 + ceil(max(B));
    end
    ang_divs = 180;
    ang_sect = (360/ang_divs);
    con = pi/180;

    for ch = 1:3    
        if ch==1
            vid = Mask;
        else
            vid = IP{1,ch};
        end
        res = zeros((R*ang_divs)+1,1);

        for t = T(1):1:T(2)
            fr = vid(:,:,t);
            Tsect = zeros(1,ang_sect);
            for div = 1:1:ang_divs
                sect = zeros(R,ang_sect);
                for r = 1:1:R
                    for sect_ang = 1:1:ang_sect
                        ang = sect_ang + ((div-1)*30);
                        x = cmx + (r*cos(ang*con));
                        y = cmy + (r*sin(ang*con));
                        sect(R+1-r,sect_ang) = fr(round(x),round(y));
                    end
                end
                Tsect = vertcat(Tsect,sect);
            end
            res = horzcat(res,Tsect);
        end

        res = res(2:size(res,1),2:size(res,2));

        %-------------------------------------------------------------------------

        res2 = zeros(1,size(res,2));

        for i = [1 24 46 68 91 113 136 158]
            res_kymo = res(((i-1)*R)+1:((i-1)*R)+R,:);
            res2 = vertcat(res2,res_kymo);
        end

        res2 = res(2:size(res2,1),:);

        if ch==1
            res2 = 65535*(1-res2);
        end

        %figure;
        %imagesc(res2); colorbar;

        %-------------------------------------------------------------

        imwrite(uint16(res2),savename,'WriteMode','append');
    end

end


%%
M = X;
range = -20:2:30;

figure;
h1 = histogram(M,range,'DisplayStyle','stairs','EdgeColor',[1 1 1],'EdgeAlpha',0,'LineWidth',1.5,'Normalization','probability');
D1 = h1.Values;
L1 = length(D1);
plot((range(1:L1)+((range(2)-range(1))/2)),(D1),'Color',[0 0.4470 0.7410],'LineWidth',3,'LineStyle','-');

set(gca,'fontsize', 18);%-------
set(gca,'FontName', 'Calibri');%-------

%%
M1 = X1;
M2 = X2;
range = -20:2:30;

figure;
hold on;
h1 = histogram(M1,range,'DisplayStyle','stairs','EdgeColor',[1 1 1],'EdgeAlpha',0,'LineWidth',1.5,'Normalization','probability');
h2 = histogram(M2,range,'DisplayStyle','stairs','EdgeColor',[1 1 1],'EdgeAlpha',0,'LineWidth',1.5,'Normalization','probability');
D1 = h1.Values;
L1 = length(D1);
D2 = h2.Values;
L2 = length(D2);
plot((range(1:L1)+((range(2)-range(1))/2)),(D1),'Color',[0 0 0],'LineWidth',3,'LineStyle','-');
D2 = h2.Values;
L1 = length(D2);
plot((range(1:L2)+((range(2)-range(1))/2)),(D2),'Color',[1 0 0],'LineWidth',3,'LineStyle','-');
set(gca,'fontsize', 18);%-------
set(gca,'FontName', 'Calibri');%-------
```
