## Supplementary material for "N-WASP-dependent branched actin polymerization attenuates B-cell receptor signaling by increasing the molecular density of receptor clusters": MATLAB scripts: TIRFvid_saveIMstack_v1.m

```
function [] = TIRFvid_saveIMstack_v1(im,filename)

len = length(filename);
if strcmp(filename(len-4:len),'.tiff') == 0
    filename = [filename '.tiff'];
end

if exist(filename) == 2
    resp = questdlg('There is already a file by that name. Would you like to overwrite it?','Overwrite Existing File?','Yes','No, cancel SAVE','Yes');
    if strcmp(resp,'Yes')==1
        delete(filename);
        imwrite(im(:,:,1),filename,'WriteMode','append');
        for i = 2:1:size(im,3)
            imwrite(im(:,:,i),filename,'WriteMode','append');
        end
    end
else
    imwrite(im(:,:,1),filename,'WriteMode','append');
    for i = 2:1:size(im,3)
        imwrite(im(:,:,i),filename,'WriteMode','append');
    end
end

end
```
