## Supplementary material for "N-WASP-dependent branched actin polymerization attenuates B-cell receptor signaling by increasing the molecular density of receptor clusters": MATLAB scripts: TIRFvid_vTrack_v1.m

```
%% TIRFvid_vTrack_v1

%Detect BCR clusters and save detection binary to file.
%Acquire BCR tracks and view tracks and track-plots.
%Quantify BCR cluster data and view plots vs time.

%% DETECT OBJECTS (LOAD VDATA FIRST!!!)

VdataN = 32;% Which cell in Vdata??
%----DO NOT EDIT PAST THIS POINT!------------------------------------------
cellData = Vdata(VdataN);
IP = openTIF_vid(cellData.fName,3);
vidR = IP{1,2};%inputs red image (BCR)
vidMask = cellData.Mask_IRM;
SFR = cellData.SFR;
D1 = size(vidMask,1);
D2 = size(vidMask,2);
sem = strel('disk',2);
cMask = 0*vidMask;
czA = zeros(1,size(cMask,3));
tn = size(vidMask,3);
MBG = zeros(1,tn);
for fr = SFR:tn
    imR = vidR(:,:,fr);
    imI = vidMask(:,:,fr);
    czA(fr) = bwarea(imI);
    R_bg = [];
    for i = 1:D1
        for j = 1:D2
            if imI(i,j)==0
                R_bg = [R_bg imR(i,j)];
            end
        end
    end
    MBG(1,fr) = median(R_bg);
    th_R = [4.1 3.9 3.7 3.5 3.3 3.1 2.9 2.7 2.5 2.3 2.1 1.9 1.7 1.5 1.3 1.1]*median(R_bg);%  <-----EDIT BCR THRESHOLDs!!!
    RMask = cell(1,size(th_R,2));
    th_R1 = th_R(1);
    imRi1 = double(imR>th_R1);
    imRi1 = imRi1.*imI;
    imRi1 = imfill(imRi1,'holes');
    imRi1 = bwareaopen(imRi1,3,4);
    se = strel('disk',1,0);
    [imlb1,n1] = bwlabel(imRi1);
    for j = 1:n1
        obj = imlb1==j;
        if sum(obj(:))<10
            imRi1 = imRi1 - obj;
            imRi1 = imRi1 + imdilate(obj,se);
        end
    end
    Mask1 = imerode(imRi1,se);
    Mask1 = imdilate(Mask1,se);
    RMask{1,1} = Mask1;
    R_Mask = Mask1*size(th_R,2);
    for i = 2:size(th_R,2)
        th_R2 = th_R(i);
        imRi2 = double(imR>th_R2);
        imRi2 = imRi2.*imI;
        imRi2 = imfill(imRi2,'holes');
        imRi2 = bwareaopen(imRi2,3,4);
        [imlb2,n1] = bwlabel(imRi2);
        for j = 1:n1
            obj = imlb2==j;
            overlap = obj.*imRi1;
            objArea = sum(obj(:));
            if sum(overlap(:)>0)||(objArea>1000)%Area upper-cutoff
                imRi2 = imRi2 - obj;
            else
                if sum(obj(:))<10
                    imRi2 = imRi2 - obj;
                    imRi2 = imRi2 + imdilate(obj,se);
                end
            end
        end
        imRi1 = imRi1 + imRi2;
        Mask2 = imerode(imRi2,se);
        Mask2 = imdilate(Mask2,se);
        RMask{1,i} = Mask2;
        R_Mask = (R_Mask + (Mask2*(size(th_R,2)+1-i)));
    end
    cMask(:,:,fr) = R_Mask>0;
end

%% SAVE RAW CLUSTER DETECTION MASK (VIDEO).

filename = cellData.fName;
filename = [filename(1:(length(filename)-4)) '_clusters-mask.tiff'];
TIRFvid_saveIMstack_v1(cMask,filename);

%% SAVE RAW CLUSTER BOUNDARIES (VIDEO).
seb = strel('disk',1);
bMask = 0*cMask;
for i = 1:1:size(cMask,3)
    bMask(:,:,i) = imdilate(cMask(:,:,i),seb) - cMask(:,:,i);
end
filename = [filename(1:(length(filename)-4)) '_cluster boundaries.tiff'];
TIRFvid_saveIMstack_v1(bMask,filename);

%% CHECK AREA VS TIME PLOT TO DETERMINE 'SWFR': THE FRAME WHEN CONTRACTION BEGINS.

figure;
plot([1:1:size(czA,2)],czA,'b.-');
xlabel('frame #');
ylabel('Area (pix)');

%% CHECK AREA VS TIME PLOT TO DETERMINE 'SWFR': THE FRAME WHEN CONTRACTION BEGINS.

figure;
plot(2*[1:1:size(czA,2)],czA,'b.-');
xlabel('Time (s)');
ylabel('Area (pix)');

%% CREATE FRAME-BY-FRAME CLUSTER ASSOCIATIONS (BASICALLY TRACK THEM).

SWFR = 115;% EDIT = the frame when contraction begins....

%----DO NOT EDIT PAST THIS POINT!------------------------------------------
DET = cell(1,1);
DETP = DET;
stop = SWFR + 61;% EDIT range (after contraction begins)
start = SWFR - 11;% EDIT range (before spreading ends)
disp(['start FR = ' num2str(start+1)]);
disp(['stop FR = ' num2str(stop-1)]);
count = 1;
for fr = stop%:-1:start
    maskFRi = uint16(cMask(:,:,fr));
    [imlbi,nci] = bwlabel(maskFRi);
    Pi = [];
    PiP = [];
    for ob = 1:1:nci
        obj_i = imlbi==ob;
        Pi(ob) = count;
        PiP(ob) = ob;
        count = count+1;
    end
    DET{fr-start+1,1} = Pi';
    DETP{fr-start+1,1} = PiP';
end
for fr = stop-1%:-1:start
    maskFRj = uint16(cMask(:,:,fr));
    [imlbj,ncj] = bwlabel(maskFRj);
    Pj = [];
    PjP = [];
    for ob = 1:1:ncj
        obj_j = imlbj==ob;
        tracked = trackObj_v1(obj_j,imlbi,1,1,3);%-----------EDIT
        if tracked==0
            Pj(ob) = count;
            PjP(ob) = ob;
            count = count+1;
        else
            Pj(ob) = Pi(tracked);
            PjP(ob) = ob;
        end
    end
    DET{fr-start+1,1} = Pj';
    DETP{fr-start+1,1} = PjP';
end
for fr = stop-2:-1:start
    maskFRk = uint16(cMask(:,:,fr));
    [imlbk,nck] = bwlabel(maskFRk);
    Pk = [];
    PkP = [];
    for ob = 1:1:nck
        obj_k = imlbk==ob;
        tracked = trackObj_v1(obj_k,imlbj,1,1,3);%-----------EDIT
        if tracked==0
            tracked = trackObj_v1(obj_k,imlbi,1,1,3);%-----------EDIT
            if tracked==0
                Pk(ob) = count;
                PkP(ob) = ob;
                count = count+1;
            else
                Pk(ob) = Pi(tracked);
                PkP(ob) = ob;
            end
        else
            Pk(ob) = Pj(tracked);
            PkP(ob) = ob;
        end
    end
    DET{fr-start+1,1} = Pk';
    DETP{fr-start+1,1} = PkP';
    Pi = Pj;
    Pj = Pk;
    imlbi = imlbj;
    imlbj = imlbk;
end
RES1 = zeros(count-1,size(DET,1));
RES2 = RES1;
for i = 1:size(DET,1)
    mat = DET{i,1};
    matP = DETP{i,1};
    for j = 1:size(mat,1)
        RES1(mat(j,1),i) = mat(j,1);
        RES2(mat(j,1),i) = matP(j,1);
    end
end

%% DELETE TRACKS SMALLER THAN A CERTAIN THRESHOLD LENGTH (IN FRAMES).

fr_thresh = 10;%---EDIT---

%----DO NOT EDIT PAST THIS POINT!------------------------------------------
RES_bin = RES1>0;
for i = 1:size(RES_bin,1)
    if sum(RES_bin(i,:))<fr_thresh
        RES2(i,:) = 0*RES2(i,:);
    end
end

figure;
hold on;
for i = 1:size(RES1,1)
    for j = 1:size(RES1,2)
        if RES1(i,j)>0
            plot(j,RES1(i,j),'r.');
        end
        if RES2(i,j)>0
            plot(j,RES1(i,j),'b.');
        end
    end
end
hold off;
xlabel(['Frame # (contraction begins at #' num2str(SWFR-start+1) ')']);
ylabel('Cluster ID');

%% Prepare corrected video Mask of clusters (dMask).

dMask = 0*cMask;
for i = 1:1:size(RES2,1)
    for j = size(RES2,2)%:-1:1
        ob = RES2(i,j);
        if ob>0
            fr = cMask(:,:,start+j-1);
            mlb = bwlabel(fr);
            obj = mlb==ob;
            dMask(:,:,start+j-1) = [dMask(:,:,start+j-1) + obj];
        end
    end
end
for i = 1:1:size(RES2,1)
    for j = size(RES2,2)-1:-1:2
        ob = RES2(i,j);
        if ob>0
            fr = cMask(:,:,start+j-1);
            mlb = bwlabel(fr);
            obj = mlb==ob;
            dMask(:,:,start+j-1) = [dMask(:,:,start+j-1) + obj];
        else
            if RES2(i,j-1)>0
                fr = cMask(:,:,start+j-2);
                mlb = bwlabel(fr);
                obj = mlb==RES2(i,j-1);
                dMask(:,:,start+j-1) = [dMask(:,:,start+j-1) + obj];
            end
        end
    end
end
for i = 1:1:size(RES2,1)
    for j = 1
        ob = RES2(i,j);
        if ob>0
            fr = cMask(:,:,start+j-1);
            mlb = bwlabel(fr);
            obj = mlb==ob;
            dMask(:,:,start+j-1) = [dMask(:,:,start+j-1) + obj];
        end
    end
end
dMask = dMask(:,:,start:stop);

%% Save dMask.

OutFolder = 'C:\Users\anbha\Desktop\Automatic tracking\DEMO\';
CellName = 'INH043_vid1_cell-03';

%----DO NOT EDIT PAST THIS POINT!------------------------------------------
filename = [OutFolder CellName '_clusters-tracked_v1.tiff'];
TIRFvid_saveIMstack_v1(uint16(dMask),filename);

%% Quantify.

QC = cell(size(RES2,1),size(RES2,2));
mbg_mat = MBG(1,start:stop);
for i = 1:1:size(RES2,1)
    for j = size(RES2,2)%:-1:1
        ob = RES2(i,j);
        bg = mbg_mat(1,j);
        if ob>0
            fr = cMask(:,:,start+j-1);
            mlb = bwlabel(fr);
            bin = mlb==ob;
            RFR = vidR(:,:,start+j-1);
            obj_ID = [CellName '__obj-' num2str(i,'%04.f')];
            Q = analObj_v1(RFR,bin,bg,obj_ID,(start+j-1),(start-SWFR-1+j));%frame correction....
            QC{i,j} = Q;
        end
    end
end
for i = 1:1:size(RES2,1)
    for j = size(RES2,2):-1:2
        ob = RES2(i,j);
        bg = mbg_mat(1,j);
        if ob>0
            fr = cMask(:,:,start+j-1);
            mlb = bwlabel(fr);
            bin = mlb==ob;
            RFR = vidR(:,:,start+j-1);
            obj_ID = [CellName '__obj-' num2str(i,'%04.f')];
            Q = analObj_v1(RFR,bin,bg,obj_ID,(start+j-1),(start-SWFR-1+j));%frame correction....
            QC{i,j} = Q;
        else
            if RES2(i,j-1)>0
                fr = cMask(:,:,start+j-2);
                mlb = bwlabel(fr);
                bin = mlb==RES2(i,j-1);
                RFR = vidR(:,:,start+j-1);
                obj_ID = [CellName '__obj-' num2str(i,'%04.f')];
                Q = analObj_v1(RFR,bin,bg,obj_ID,(start+j-1),(start-SWFR-1+j));%frame correction....
                QC{i,j} = Q;
            end
        end
    end
end
for i = 1:1:size(RES2,1)
    for j = 1
        ob = RES2(i,j);
        bg = mbg_mat(1,j);
        if ob>0
            fr = cMask(:,:,start+j-1);
            mlb = bwlabel(fr);
            bin = mlb==ob;
            RFR = vidR(:,:,start+j-1);
            obj_ID = [CellName '__obj-' num2str(i,'%04.f')];
            Q = analObj_v1(RFR,bin,bg,obj_ID,(start+j-1),(start-SWFR-1+j));%frame correction....
            QC{i,j} = Q;
        end
    end
end
for i = size(RES2,1):-1:1
    RES_row = RES2(i,:);
    if sum(RES_row)==0
        QC(i,:) = [];
    end
end
QC(:,size(QC,2)) = [];
QC(:,1) = [];

QCN = QC;
for i = 1:size(QC,1)
    for j = 1:size(QC,2)
        if ~isempty(QC{i,j})
            Q = QCN{i,j};
            bg = Q.BG;
            MFI_n = Q.MFI/bg;
            Peak_n = Q.Peak/bg;
            Q.MFIN = MFI_n;
            Q.PeakN = Peak_n;
            QCN{i,j} = Q;
        end
    end
end            

%% Plot cluster tracks (all in 1).

%Xcell = cell(1,size(QC,1)); Ycell = cell(1,size(QC,1));
figure;
hold on;
for i = 1:size(QC,1)
    X = [];
    Y = [];
    for j = 1:size(QC,2)
        if ~isempty(QC{i,j})
            Q = QC{i,j};
            X = [X Q.CX];
            Y = [Y Q.CY];
        end
    end
    Z = Y;
    Y = X;
    X = Z;
    %figure;%
    %hold on;%
    plot(X,Y,'k.-');
    plot(X(1),Y(1),'g.');
    plot(X(length(X)),Y(length(Y)),'r.');
    %hold off;%
    %Xcell{1,i} = X;
    %Ycell{1,i} = Y;
    %xlim([1 120]);%
    %ylim([1 120]);%
end
hold off;
xlim([1 D2]);
ylim([1 D1]);
axis ij;
%axis square;


%% Plot data (one field of choice).

Dfield = 'Peak';%------WHAT TO PLOT???------------------

%----DO NOT EDIT PAST THIS POINT!------------------------------------------
mbg = MBG(1,(start+1):(stop-1));
figure;
hold on;
for i = 1:size(QC,1)
    T = [];
    Y = [];
    for j = 1:size(QC,2)
        if ~isempty(QC{i,j})
            Q = QC{i,j};
            VAL =[];
            eval(['VAL = Q.' Dfield ';']);
            Y = [Y VAL];
            T = [T Q.cFR];
        end
    end
    plot(2*T,Y,'k.-');
end
plot((-2*(SWFR-start-1):2:2*(stop-SWFR-1)),mbg,'r.');
hold off;
ylabel(['BCR cluster ' Dfield]);
xlabel('Time (s) [contraction begins at 0]');
xlim([-2*(SWFR-start-1) 2*(stop-SWFR-1)]);
ylim([0 1000]);%---EDIT---


%% Plot data (all fields).

Fields = cell(1,5);
Fields{1,1} = 'ID';
Fields{1,2} = 'TFI';
Fields{1,3} = 'MFI';
Fields{1,4} = 'Area';
Fields{1,5} = 'PbM';
Fields{1,6} = 'Peak';%------WHAT TO PLOT???------------------

%----DO NOT EDIT PAST THIS POINT!------------------------------------------

mbg = MBG(1,(start+1):(stop-1));
figure;
for k = 1:size(Fields,2)
    Dfield = Fields{1,k};
    subplot(2,3,k);
    hold on;
    for i = 1:size(QC,1)
        T = [];
        Y = [];
        for j = 1:size(QC,2)
            if ~isempty(QC{i,j})
                Q = QC{i,j};
                VAL =[];
                if strcmp(Dfield,'ID')
                    VAL = i;
                else
                    eval(['VAL = Q.' Dfield ';']);
                end
                Y = [Y VAL];
                T = [T Q.cFR];
            end
        end
        plot(2*T,Y,'k.-');
    end
    if (k==3)||(k==6)
        plot((-2*(SWFR-start-1):2:2*(stop-SWFR-1)),mbg,'r.');
    end
    hold off;
    ylabel(['BCR cluster ' Dfield]);
    xlabel('Time (s) [contraction begins at 0]');
    xlim([-2*(SWFR-start-1) 2*(stop-SWFR-1)]);
end


%% Plot data 3 (plot all fields after isolating clusters spanning contraction).

QCD = QC;
for i = size(QC,1):-1:1
    TC = [];
    for j = 1:1:size(QC,2)
        if ~isempty(QC{i,j})
            Q = QC{i,j};
            TC = [TC Q.cFR];
        end
    end
    if (min(TC)>0)||(max(TC)<5)
        QCD(i,:) = [];
    end
end

Fields = cell(1,5);
Fields{1,1} = 'ID';
Fields{1,2} = 'TFI';
Fields{1,3} = 'MFI';
Fields{1,4} = 'Area';
Fields{1,5} = 'PbM';
Fields{1,6} = 'Peak';

mbg = MBG(1,(start+1):(stop-1));
figure;
for k = 1:size(Fields,2)
    Dfield = Fields{1,k};
    subplot(2,3,k);
    hold on;
    for i = 1:size(QCD,1)
        T = [];
        Y = [];
        for j = 1:size(QCD,2)
            if ~isempty(QCD{i,j})
                Q = QCD{i,j};
                VAL =[];
                if strcmp(Dfield,'ID')
                    VAL = i;
                else
                    eval(['VAL = Q.' Dfield ';']);
                end
                Y = [Y VAL];
                T = [T Q.cFR];
            end
        end
        plot(2*T,Y,'k.-');
    end
    if (k==3)||(k==6)
        plot((-2*(SWFR-start-1):2:2*(stop-SWFR-1)),mbg,'r.');
    end
    hold off;
    ylabel(['BCR cluster ' Dfield]);
    xlabel('Time (s) [contraction begins at 0]');
    xlim([-2*(SWFR-start-1) 2*(stop-SWFR-1)]);
end

%% Plot data 4 (plot shift in MFI and Peak of clusters during contraction).

MFImat = zeros(size(QCD,1),2);
Peakmat = zeros(size(QCD,1),2);
mbg = MBG(1,(start+1):(stop-1));
mbgt1 = mbg(1,(SWFR-start));
for i = 1:1:size(QCD,1)
    Qt1 = QCD{i,(SWFR-start)};
    mfit1 = Qt1.MFI;
    maxt1 = Qt1.Peak;
    tt1 = 0;
    for j = size(QCD,2):-1:1
        if ~isempty(QCD{i,j})
            break;
        end
    end
    Qt2 = QCD{i,j};
    mbgt2 = mbg(1,j);
    mfit2 = (Qt2.MFI)*(mbgt1/mbgt2);
    maxt2 = (Qt2.Peak)*(mbgt1/mbgt2);
    tt2 = Qt2.cFR;
    MFImat(i,1) = mfit1;
    MFImat(i,2) = mfit2;
    Peakmat(i,1) = maxt1;
    Peakmat(i,2) = maxt2;
end

figure;%plots MFI
hold on;
for i = 1:1:size(QCD,1)
    MFIi = MFImat(i,:);
    plot([1 2],MFIi,'r-');
    plot([1 2],MFIi,'k.','MarkerSize',10);
end
ylabel(['BCR cluster MFI']);
xlim([0 3]);
ylim([0 1000]);
xticks([]);
[hyp,pval] = ttest(MFImat(:,1),MFImat(:,2));
text(0.5,100,['p value (paired ttest) = ' num2str(pval)]);
hold off;

figure;%plots Peak
hold on;
for i = 1:1:size(QCD,1)
    MAXi = MFImat(i,:);
    plot([1 2],MAXi,'r-');
    plot([1 2],MAXi,'k.','MarkerSize',10);
end
ylabel(['BCR cluster Peak']);
xlim([0 3]);
ylim([0 1000]);
xticks([]);
[hyp,pval] = ttest(Peakmat(:,1),Peakmat(:,2));
text(0.5,100,['p value (paired ttest) = ' num2str(pval)]);
hold off;
```
