## Supplementary material for "N-WASP-dependent branched actin polymerization attenuates B-cell receptor signaling by increasing the molecular density of receptor clusters": MATLAB scripts: trackObj_v1.m

```
function [REF_OBJ] = trackObj_v1(obj,ref,obj_dil,ref_dil,min_ovrlp)

%obj = X;
%ref = Y;
%ref_dil = 2;
%obj_dil = 2;
%min_ovrlp = 1;

if obj_dil>0
    DOB = imdilate(obj,strel('disk',obj_dil));
else
    DOB = obj;
end

List = [];
Lcount = 1;
[reflb,N] = bwlabel(ref);
for i = 1:N
    P = reflb==i;
    if ref_dil>0
        DP = imdilate(P,strel('disk',ref_dil));
    else
        DP = P;
    end
    OL = DP.*DOB;
    if sum(OL(:))>=min_ovrlp
        List(Lcount,1) = i;
        List(Lcount,2) = sum(OL(:));
        Lcount = Lcount + 1;
    end
end

if isempty(List)
    REF_OBJ = 0;
else
    List = sortrows(List,2,'descend');
    REF_OBJ = List(1,1);
end

end
```
